# Enabling Atomistic Modeling and Simulation of Complex Curved Cellular Membranes with xMAS Builder

**DOI:** 10.1101/2025.01.14.632907

**Authors:** Noah Trebesch, Emad Tajkhorshid

## Abstract

As more powerful high performance computing resources are becoming available, there is a new opportunity to bring the unique capabilities of molecular dynamics (MD) simulations to cell-scale systems. Membranes are ubiquitous within cells and are responsible for a diverse set of essential biological functions, but building atomistic models of cell-scale membranes for MD simulations is immensely challenging because of their vast sizes, complex geometries, and complex compositions. To meet this challenge, we have developed xMAS Builder (E**x**perimentally-Derived **M**embranes of **A**rbitrary **S**hape Builder), which is designed to take experimental lipidomics and structural (e.g., electron microscopy and tomography) data as input and use them to build MD-ready models of cellular membrane systems. To test xMAS Builder’s capabilities, we have used it to build two models (one ∼12.0 million atoms and the other ∼11.6 million atoms) of a test system with a representative complex lipid composition and geometry. The two models, which differed only in their lipid packing densities, both maintained their membrane integrity during an extended MD simulation (250 ns and 386 ns), but their highly divergent relaxation dynamics indicate that the proper packing density of curved membranes is determined by leaflet volume rather than surface area. These results suggest that xMAS Builder’s algorithms produce high quality models and that simulation of these models will provide profound biophysical insights into the behavior of cellular membranes.

## 1 Introduction

Molecular dynamics (MD) is a powerful computational technique whose basic purpose is to elucidate the fundamental structural underpinnings of biological function through high spatial and temporal resolution simulations of the dynamics of biological systems. At present, MD has most commonly been used to study a single protein or protein complex at a time, but MD simulations have also successfully been applied to larger and more complex biological systems, including several viruses,^1–10^ a chromatophore,^11^ cellular cytoplasm,^12^ a gene locus,^13^ and even a minimal cell.^14, 15^ As increasingly powerful high performance computing systems become available, there is a new opportunity to bring the unique capabilities of MD to new classes of cell-scale biological systems.

Cell-scale membranes are prime targets for detailed characterization with MD because they are a ubiquitous part of living systems, and they play an essential role in biological function, acting as semipermeable barriers that separate and protect cells and organelles from their surroundings. Within the cell, membranes adopt a variety of highly complex shapes, captured in detail by electron microscopy (EM) and electron tomography (ET) experiments,^16–18^ and they are also highly complex in terms of their lipid and protein compositions. Combined with their immense scale, these features of cell-scale membranes pose a grand challenge to atomistic modeling, requiring a significant investment in the development of new modeling tools and techniques.

To meet this challenge, we have developed xMAS Builder (E**x**perimentally-Derived **M**embranes of **A**rbitrary **S**hape Builder), a collection of scalable modeling algorithms designed to generate MD-ready atomistic models of cellular membrane structures. Starting with a triangle mesh derived directly from EM or ET data, xMAS Builder creates a membrane model by filling the arbitrarily complex mesh shape with an arbitrarily complex lipid composition derived from lipidomics data (Fig. 1). To ensure the final membrane model can be stably simulated with MD, xMAS Builder employs a series of modeling approaches to determine the proper thickness and packing density of the membrane, to evenly distribute the lipids over the surfaces of the membrane, to resolve complex steric clashes between lipids placed in the model, and to solvate and equilibrate the model.

**Figure 1:**
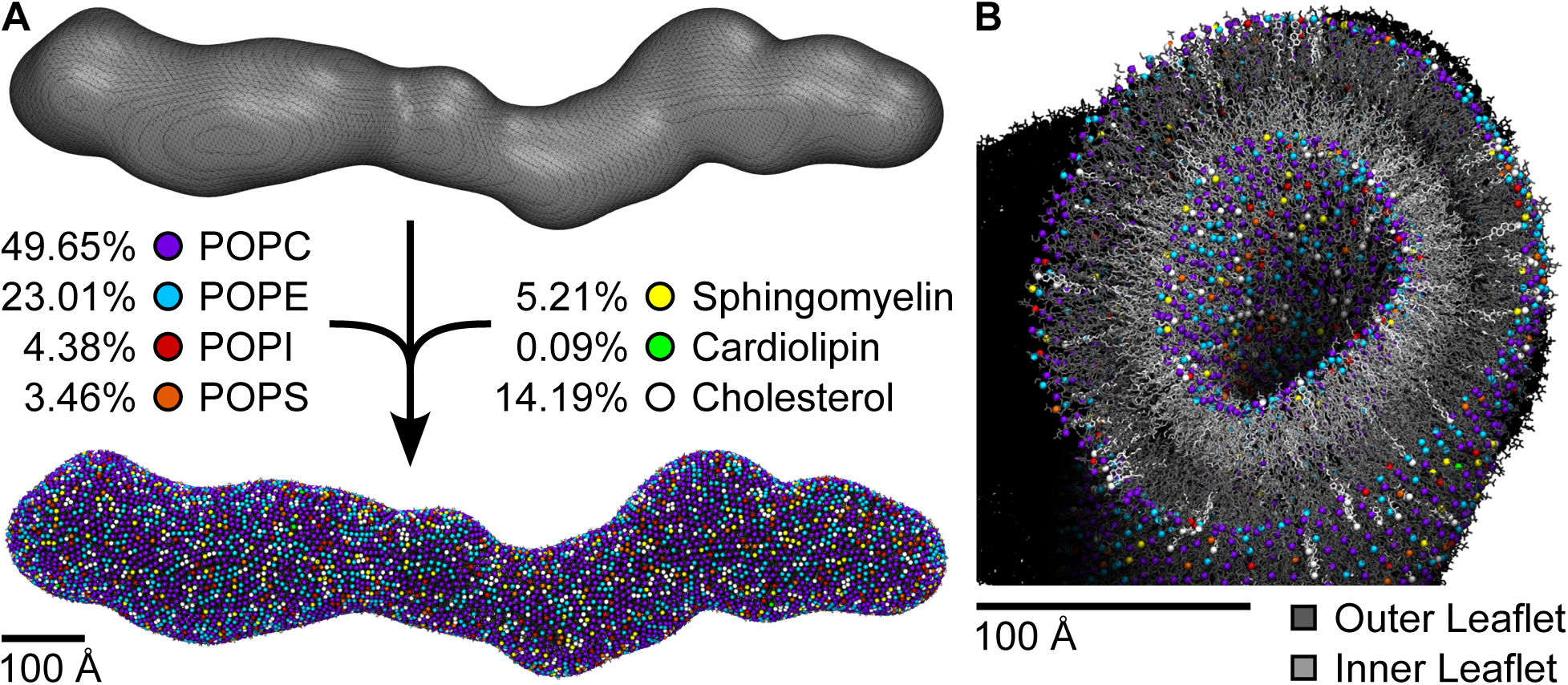
Application of xMAS Builder to a test membrane system. **(A)** xMAS Builder uses as input a mesh representing the surface of the membrane and a target lipid composition and turns them into an atomistic model that can be stably simulated using MD. In the application highlighted here, the input mesh is synthetic and is meant to capture the complex curvature of natural cellular membranes while being much smaller than them to make the development of xMAS Builder tractable. The input lipid composition is derived from an experimentally-determined lipid composition of the ER and is meant to be representative of the compositional complexity of cellular membranes. **(B)** Cutaway molecular rendering of the model generated by xMAS Builder viewed down its long axis. For each lipid, one atom near the membrane surfaces is represented as a sphere (the oxygen atom for cholesterol, the central linking carbon atom for cardiolipin, and the phosphorus atom for POPC, POPE, POPI, POPS, and sphingomyelin). In (A), the spheres have double the radius to better highlight the positions and identities of individual lipids.

xMAS Builder is not unique in its goal to generate MD-ready models of membrane systems. Many tools have been developed to generate membrane models with simple planar or spherical shapes, including CHARMM-GUI,^19–24^ insane,^25^ PACKMOL-Memgen,^26^ and others. Other tools, namely PACKMOL^27^ and BUMPy,^28^ are designed to generate more complex membrane geometries through user-specified combinations of a handful of simple parametric shapes. cellPACK^29^ is capable of packing lipids and proteins into mesoscale membrane models, but the significant challenge of adapting these models for use in MD simulations has not been addressed. Ultimately, only two tools, LipidWrapper^30^ and TS2CG,^31^ are capable of generating MD-ready membrane models with arbitrarily complex shapes, and we will discuss how they compare to xMAS Builder after describing its approach in detail.

xMAS Builder’s modeling approaches have all been designed to handle cell-scale membrane models, but it was impractical to rapidly develop, troubleshoot, and test the efficacy of new approaches using a full cell-scale system. Instead, we used a smaller (∼1,124 Å by ∼260 Å by ∼268 Å) synthetic test system with a realistically complex membrane shape, and we filled it using a lipid composition of the endoplasmic reticulum (ER) derived from experimental data^32^ (Fig. 1). We ultimately produced two models of the test system, one containing ∼12.0 million atoms and the other ∼11.6 million atoms, that differed only in their lipid packing densities, and we evaluated the stability and quality of the models using long-term (250 ns and 386 ns, respectively) MD simulations. Here, we describe each of the new modeling approaches we developed for xMAS Builder, we report the results of applying them to the test system, and we discuss the results from the long-term MD simulations of the two versions of the model.

## 2 Results

xMAS Builder’s approach to building models of cellular membranes has four major modeling stages:

1. Determine the lipid packing density and thickness of the membrane.
2. Place the lipids within the membrane.
3. Detect and eliminate complex steric clashes between lipids.
4. Solvate and equilibrate the membrane model.

For each of these modeling stages, we have developed new algorithms designed to tackle the unique challenges associated with building cellular membrane models, and, whenever appropriate, we have also applied existing algorithms in new ways. xMAS Builder is implemented as a series of Python 3 and VMD^33^ TCL scripts. Several modeling steps involve running MD simulations with NAMD,^34^ and we also make use of several standard triangle mesh algorithms implemented in MeshLab.^35^

In the rest of this section, we describe what each of xMAS Builder’s modeling algorithms do and why they are designed and/or applied in the way that they are. We have used xMAS Builder to generate two models of a test system, and as we introduce each modeling algorithm, we also discuss its application to one of these models. Later, we introduce the other, earlier version of the model, discuss the application of xMAS Builder’s modeling approaches to it, and compare it to the final model.

### 2.1 Determining the Lipid Packing Density and Membrane Thickness

When building a model with xMAS Builder, the user starts by providing two principal inputs: the desired lipid composition of the membrane model and a triangle mesh representing the shape of the membrane structure that is being modeled. This triangle mesh may be obtained from experimental sources (e.g., EM or ET) or generated synthetically, depending on the desired application. Together, these two inputs give the ultimate membrane model its defining characteristics, one of which is the number of lipid molecules present in the membrane model. Inaccurately estimating the number of lipids needed to fill a curved membrane with a complex lipid composition results in membrane defects when the model is simulated (see Sec. 2.6), so xMAS Builder’s protocol is designed to ensure the accuracy of this estimate.

#### 2.1.1 Approach

To start, an atomistic model of a planar membrane patch large enough to have a composition that is representative of the target lipid composition of the final model is prepared using CHARMM-GUI. It is then equilibrated with MD using the same simulation parameters that will be used in the final simulation of the membrane model. The point of this simulation is to determine the bulk properties of a membrane patch with the desired lipid composition, particularly its equilibrated volume and thickness.

Next, using the original mesh input and the measured thickness of the membrane patch, xMAS Builder generates meshes representing the inner and outer surfaces of the membrane as well as a surface that passes through the midplane of the membrane. This is done by sliding the original mesh vertices the appropriate distance along the vertex normal vectors while maintaining the original connectivity between the mesh vertices. A standard mesh algorithm (as implemented by the “Compute Geometric Measures” filter in MeshLab) is used to measure the volume enclosed by each of the generated meshes, and these volumes are in turn used to estimate the volume occupied by the inner and outer leaflets of the membrane model. Using these volumes, the measured volume of the membrane patch, and the input lipid composition, the number of lipids of each type needed to fill the inner and outer leaflets is estimated.

#### 2.1.2 Application

The triangle mesh and lipid composition of the test membrane system being modeled here have been previously introduced (Fig. 1). Due to the small percentage of cardiolipin in the lipid composition, we generated a large membrane patch composed of 1,000 lipids per leaflet (Fig. 2A). We equilibrated the membrane patch for 20 ns, and the thickness and volume of the membrane stabilized after ∼10 ns to ∼43.2 Å and ∼2,267,024 Å^3^, respectively (Fig. 2B-C). After xMAS Builder generated the meshes representing the outer and inner surfaces and midplane of the membrane model, it estimated the volume occupied by the outer and inner leaflets to be ∼11,057,386 Å^3^ and ∼7,192,268 Å^3^, respectively. From these numbers, xMAS Builder calculated that 9,755 and 6,345 lipids were needed to fill the outer and inner leaflets (Fig. 2D).

**Figure 2:**
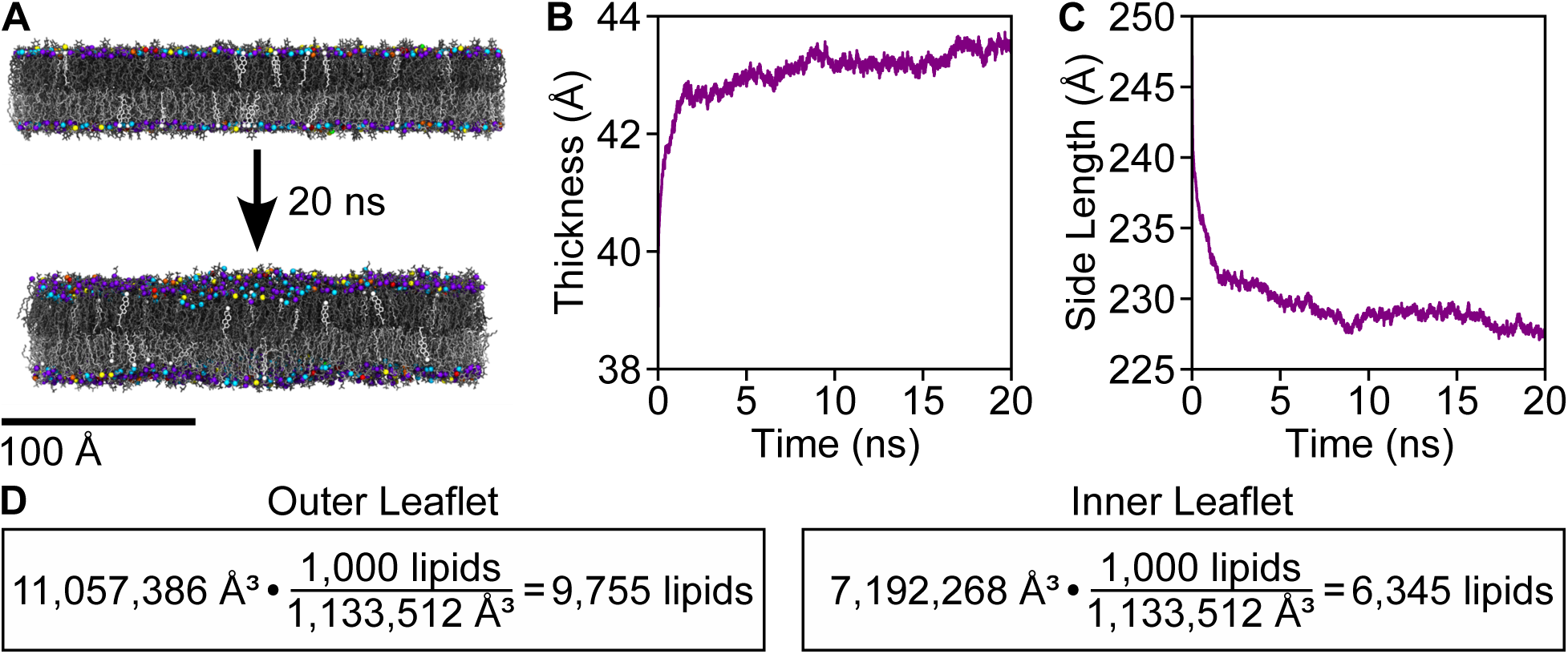
Determining the lipid packing density and thickness of the membrane. **(A)** Molecular renderings of a planar membrane patch with the target lipid composition before and after 20 ns of simulation. The point of this simulation was to determine the physical parameters to use for the membrane model, which change depending on the target lipid composition. The same colors and molecular representations established in Fig. 1 are also used here. **(B)** Plot of the thickness of the membrane as a function of time during the simulation. This measurement determines how far apart lipids in the membrane’s two leaflets are placed when the model is initially built. **(C)** Plot of the side length of the membrane patch as a function of time during the simulation. Together with the thickness measurement, this measurement determines the volume of the patch, which in turn determines the lipid packing density used for the membrane model. **(D)** Calculations of the number of lipids used to fill the inner and outer leaflets in the model, determined by multiplying the volume of the inner and outer leaflets of the membrane by the lipid packing density of the equilibrated membrane patch.

### 2.2 Optimizing the Initial Placement of the Lipids

Once the number of lipids needed to fill the inner and outer leaflets has been calculated, xMAS Builder must place the lipids within the leaflets. The most straightforward way to do this is to use the positions of vertices from the meshes representing the inner and outer surfaces of the membrane. This approach requires that the meshes have at least as many vertices as there are lipids in the model, which can be achieved using standard mesh upsampling algorithms if needed. Unfortunately, this direct approach also introduces some consequential artifacts into the membrane model.

One problem is that mesh vertices are not uniformly distributed over the surface of a mesh, so using this approach results in an uneven initial packing of the placed lipids. Additionally, it is difficult to control the exact number of vertices within a mesh, so meshes with slightly more vertices than there are lipids have to be used in practice. As such, not every vertex has a lipid placed at its position, creating small voids that compound the uneven distribution of the placed lipids. Given enough equilibration time, these artifacts would dissipate. However, because of the large scale of the models xMAS Builder is designed to build, it is essential to minimize the amount of equilibration that must be performed on the full membrane model before it can start producing meaningful results. Accordingly, we have designed a new scalable and computationally efficient approach to optimize the initial placement of the lipids with xMAS Builder.

#### 2.2.1 Approach

In xMAS Builder’s approach, a single particle is used to represent each lipid in the system (Fig. 3A). The particles are initially placed on the vertices of the meshes representing the inner and outer surfaces of the membrane, meaning they are not initially evenly distributed on the surfaces of the meshes. To optimize their positions, they are simulated using MD while restrained to the inner and outer surfaces of the membrane model. Basing xMAS Builder’s approach on an MD simulation allows it to harness the established parallel performance of NAMD to optimize the initial positions of millions of lipids at once, and the small particle count of the simulation gives it a highly economical computational cost.

**Figure 3:**
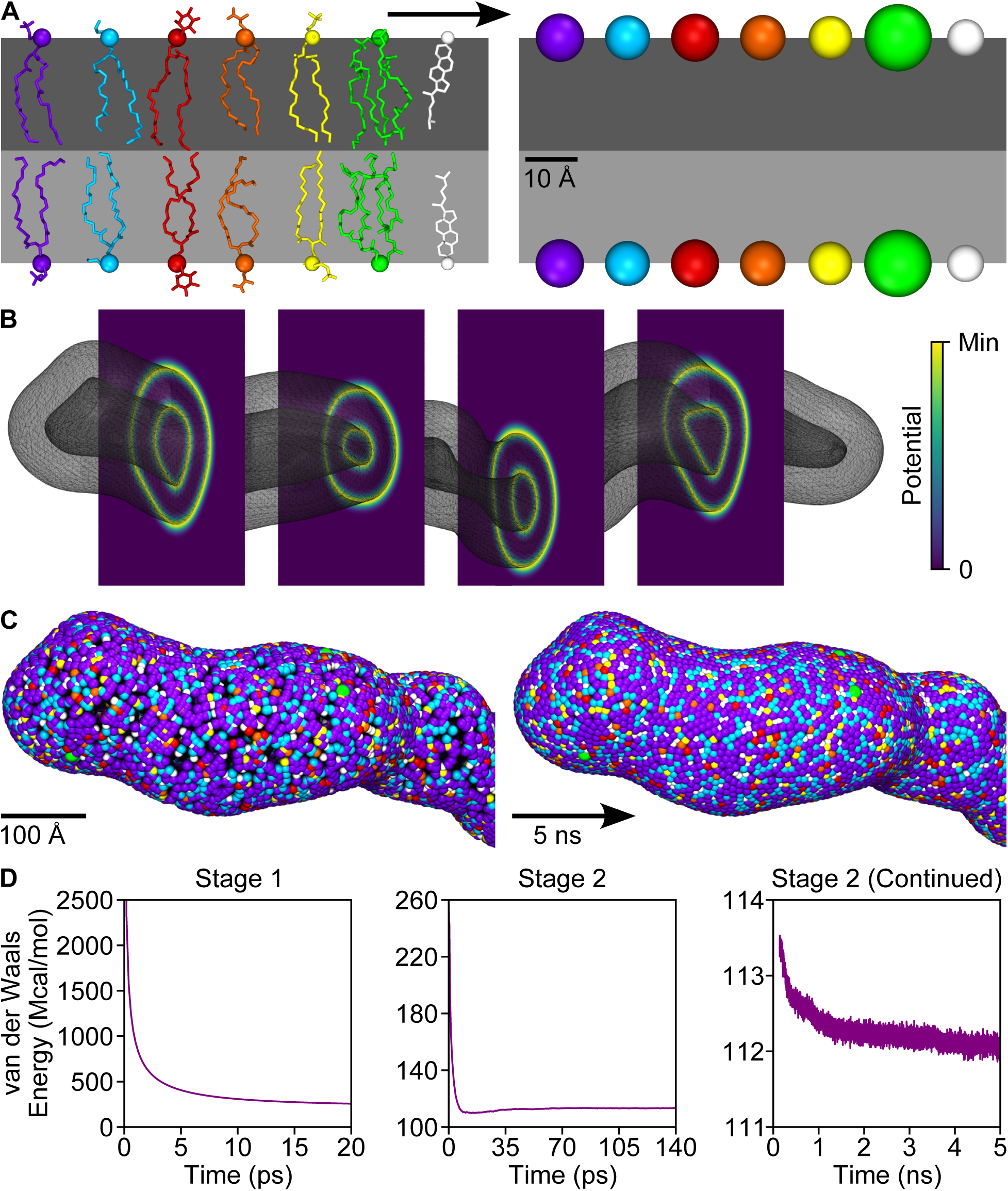
Optimizing the initial placement of the lipids. **(A)** The positions of the lipids is optimized using an MD simulation in which each lipid is represented as a single uncharged particle. The mass of the particle is determined by the mass of the lipid, and its radius is determined by the area-per-lipid value reported by CHARMM-GUI. The same colors and molecular representations established in Fig. 1 are also used here. **(B)** Visualization of the potential used by the Grid Forces feature of NAMD to restrain the lipid particles to the surfaces of the membrane during the simulation. The force applied by Grid Forces is proportional to the negative gradient of the user-specified potential, and the potential in this application has the functional form of a truncated Gaussian centered on the meshes representing the membrane surfaces. **(C)** Distribution of the lipid particles before and after the simulation. The initial positions of the lipid particles are determined by the positions of a random selection of vertices from the meshes representing the surfaces of the membrane. Large gaps can be seen between the particles at the start of the simulation, but the particles evenly distribute themselves over the membrane surface over the course of the simulation. **(D)** Plots of the total particle interaction energy during the simulation, which is used to determine when the distribution of the particles has stabilized. The simulation occurs in two stages. The first stage (20 ps) uses the velocity quenching feature of NAMD to eliminate large overlaps between particles while preventing them from moving too quickly and destabilizing the simulation. The second stage (5 ns) proceeds using standard MD.

In the simulation, the particles are designed to interact with one another solely through Lennard-Jones potentials (as specified by the CHARMM36m force field^36–39)^, and different parameters (*∈*_i_ and 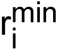 below) are used for each type of lipid in the model.

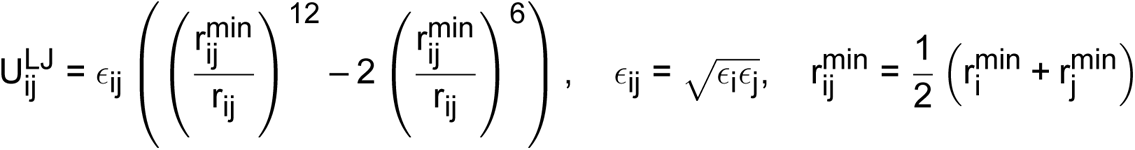

The chosen parameters are designed to ensure that the particles constantly experience van der Waals repulsion and to ensure that the space occupied by the particles is proportional to the space occupied by the lipids they represent. As such, all of the particles are given the same value of *∈*_i_ (empirically set at 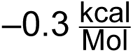), and 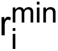 is set for each particle such that it is twice the radius of a circle with an area equal to the pertinent area-per-lipid value reported by CHARMM-GUI.

The particles are restrained to the inner and outer surfaces of the membrane using the Grid Forces^40^ feature of NAMD. In Grid Forces, an energetic potential field is created over a region of a model by assigning potential values to points on a regular three dimensional grid. Between grid points, the potential is determined using an interpolation scheme, and the force applied to each particle is determined by calculating the negative gradient of the potential multiplied by a scaling factor. To use Grid Forces, xMAS Builder generates potential grids that represent the shapes of the inner and outer surfaces of the membrane (Fig. 3B). Specifically, xMAS Builder does this by creating an empty grid over each mesh, calculating the minimum distances between each grid point and the mesh, applying a truncated Gaussian function to each of those distances, and assigning the resulting values to the original respective grid points.

With the particles representing the lipids placed and Grid Forces active, the MD simulation can begin. Due to the strength of the interaction forces between the particles, the poor initial placement of the particles makes standard MD unstable for the initial configuration of the system. As such, xMAS Builder runs the simulation at a temperature of 100 K, and standard MD is preceded by a short MD simulation with the velocity quenching feature of NAMD active, which simply prevents particles from moving more than a set distance within a time step. MD with velocity quenching is run until the interaction energy between the particles stabilizes, and standard MD is then also run until the interaction energy between the particles re-stabilizes.

After both stages of the simulation is complete, the particles must be replaced with atomistic lipids. To determine the orientations to use for the atomistic lipids, the simulation particles are separated into two groups and treated as two point clouds (one for the inner surface and one for the outer surface), and a standard mesh algorithm (as implemented by the “Compute normals for point sets” filter in MeshLab) is applied. The particles are then replaced with atomistic lipids from a library of lipid conformations derived from CHARMM-GUI, which produces a fully atomistic model of the membrane system.

#### 2.2.2 Application

After xMAS Builder generated the Grid Forces grids and placed the particles representing the lipids (Fig. 3B-C), MD with velocity quenching was performed for 20 ps, causing the interaction energy between the particles to drop from an initial value of 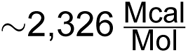 to a final value of 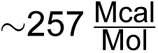 (Fig. 3D). Following MD with velocity quenching, standard MD was performed for 5 ns, during which time the interaction energy dropped to a stable value of 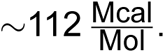 As intended, this simulation produced an even distribution of the lipid particles over the surfaces of the membrane model (Fig. 3C).

### 2.3 Fixing Complex Steric Clashes Among the Lipids

After the simulation to optimize the initial placement of the lipids, xMAS Builder uses the positions of the particles to generate a fully atomistic model of the membrane system. However, this model is not yet suitable for MD simulation because of the existence of complex steric clashes between the lipids that make up the model, namely ring piercings and lipid entanglements. A ring piercing occurs when a bond from one lipid passes through the center of a ring in another lipid, and a lipid entanglement occurs when three sets of bonded atoms are locked together in the same region of space (Fig. 4). xMAS Builder must fix these clashes before an MD simulation can be run because they do not occur in natural membrane systems, they cause MD simulations to crash, and they cannot be eliminated by applying standard MD minimization.

**Figure 4:**
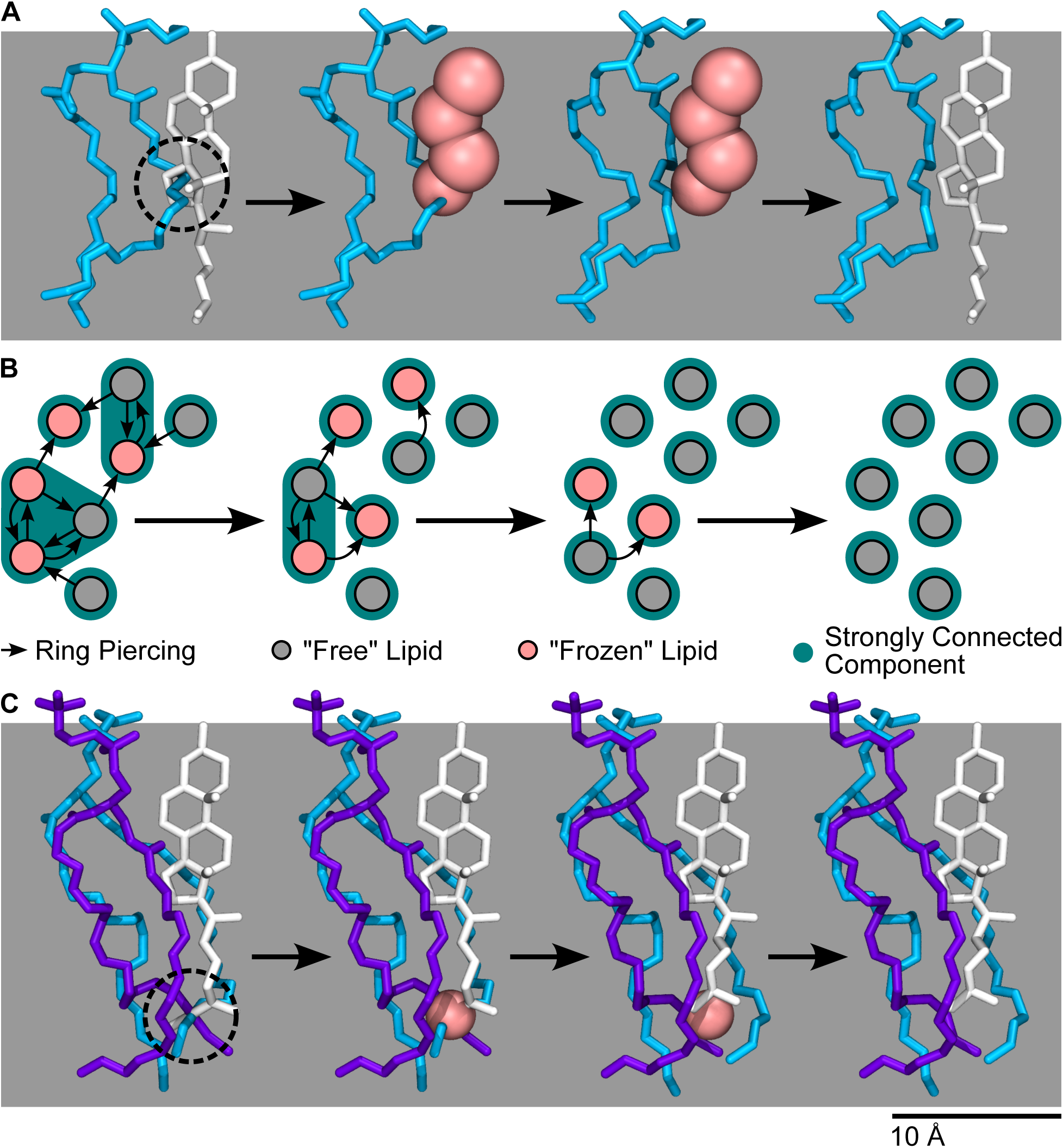
Fixing complex steric clashes between lipids. **(A)** Strategy to fix ring piercings. In the initial configuration, a POPE lipid (cyan) pierces the ring of a cholesterol lipid (white). To fix this piercing, the cholesterol lipid is removed from the membrane, and a particle (pink) representing each of its rings is added to the membrane. Then, 500 steps of minimization are performed in NAMD while keeping the positions of the ring particles fixed, which causes the POPE tail to no longer overlap with the particles. The particles are then removed from the membrane, the cholesterol lipid is added back with its original conformation, and the new conformation of the POPE lipid no longer pierces the ring of cholesterol. Because their conformation does not change during this process, we refer to the ringed lipids replaced by particles as “frozen,” and we refer to the rest of the lipids as “free.” **(B)** Strategy to handle networks of ring piercings. xMAS Builder represents networks of ring piercings as directed graphs, with nodes representing lipids and with edges directed from the piercing lipid to the pierced lipid. Within the directed graphs, xMAS Builder identifies the strongly connected components and “freezes” all but one of the lipids within the component. The lipid chosen to be “free” is the one that is pierced by the greatest number of lipids and that is not also pierced by any lipid from outside of the strongly connected component. With the “frozen” and “free” lipids selected, xMAS Builder applies the approach from (A), and networks of ring piercings are searched for again in the resulting system. These steps are applied iteratively until no ring piercings are detected in the system. **(C)** Strategy to correct lipid entanglements. In the initial configuration, bonds from a cholesterol lipid (white), a POPC lipid (purple), and a POPE lipid (cyan) are unnaturally extended and centered on the same region in space, creating a lipid entanglement. To correct this issue, xMAS Builder places a particle (pink) at the center of the entanglement and holds its position fixed during 500 steps of minimization in NAMD. The presence of the particle disrupts the energetic favorability of the entangled conformation, allowing the lipids’ atoms to move away from each other and the bonds to return to their natural lengths during minimization. At the end of this process, the particle is removed.

Complex lipid clashes are introduced into the membrane model because xMAS Builder places individual lipids blindly (i.e., without regard for the placement of other lipids). It is possible, of course, to place lipids using a non-blind approach (e.g., as in CHARMM-GUI’s^19–24^ approach), but such approaches do not scale to cell-sized membrane systems. Moreover, existing approaches to fix ring piercings sometimes fail, forcing the user to fix the remaining problems manually or rebuild the entire system from scratch. Occasional failure is fine for smaller membrane models, but it is not a viable option for cell-scale systems where the number of ring piercings is orders of magnitude larger.

Standard MD minimization is able to fix simple atomic clashes present in newly built models, and it is designed to efficiently harness the parallel computing power of MD engines. However, it is not able to correct ring piercings and lipid entanglements because, in both of these molecular configurations, opposing atomic interactions counterbalance each other. Specifically, steric clashes between the molecular groups in the system work to push the groups apart, while covalent bonds between some groups work to pull them back together. As such, to disrupt these configurations, the interaction energies between the groups need to temporarily increase through worsened steric clashes and/or lengthened covalent bonds, which minimization cannot achieve on its own because it is designed to exclusively lower interaction energies. In xMAS Builder, we have therefore developed a customized approach to fix these complex lipid clashes.

#### 2.3.1 Approach

Before ring piercings can be corrected, they must first be detected. In xMAS Builder’s approach, rings in lipids are decomposed into triangles, nearby bonds from other lipids are treated as line segments, and simple geometric tests (modified from^41^) are applied to determine if any of the line segments intersect any of the triangles. After ring piercings have been detected, they are corrected by removing lipids with pierced rings from the system, replacing each pierced ring with a large particle, holding the position of the particle fixed, and applying 500 steps of minimization to the full system (Fig. 4A). The presence of the particle representing the ring makes it energetically unfavorable for atoms to intersect with the particle, allowing minimization to gently push them away from the ring. After minimization, the particles are replaced with the original atomistic lipids. Because the conformations of the lipids replaced with particles do not change throughout this process while the conformations of the other lipids do, we refer to the replaced lipids as “frozen,” and the rest of the lipids as “free.”

This process is complicated by the existence of networks of ring piercings (i.e., collections of ringed lipids piercing each other in a cyclical manner). The simplest example of a network of ring piercings involves two lipids with two interlocked rings. If the rings of these two lipids were both replaced with particles, the subsequent minimization would accomplish nothing because the position of both particles is held fixed, and none of the atoms involved in the original piercings would be present during minimization. To handle this situation, xMAS Builder represents all ring piercings detected within a system as a directed graph, finds the connected components within the graphs (which represent the networks), finds the strongly connected components within the graph’s connected components, “freezes” all but one lipid within each strongly connected component, and applies minimization (Fig. 4B). This approach breaks networks up into multiple pieces, which are often themselves smaller networks. As such, the procedure to detect and break up networks must be applied iteratively until xMAS Builder no longer detects any ring piercings in the system.

As with ring piercings, lipid entanglements must first be detected before they can be corrected. When minimization is applied to the whole model to correct ring piercings, the bonds involved in an entanglement become highly extended, and they center themselves on the same region in space (Fig. 4C), making them straightforward to detect. Once found, xMAS Builder places a large particle at the center of each entanglement, and the position of the particle is held fixed during 500 steps of minimization. The presence of the particle creates a steric clash between it and the atoms involved in the entanglement, destabilizing the entanglement and allowing minimization to move the atoms involved away from the location of the fixed particle. After applying minimization, the large particles are removed from the system.

#### 2.3.2 Application

The test system is composed of 16,100 lipids, and 2,990 of those lipids contain rings (i.e., cholesterol and POPI). xMAS Builder took two iterations of 500 steps of minimization to resolve all of the ring piercings in the test system. Specifically, it detected 805 ring piercings in the initial system, then 12 after the first minimization iteration, then 0 after the second. For the first iteration, it selected 437 lipids to “freeze,” then it selected 7 to “freeze” for the second iteration. After correcting the ring piercings, xMAS Builder detected that there were 3 lipid entanglements in the system, which it then resolved in 500 steps of minimization. Thus, xMAS Builder corrected all of the complex lipid clashes in the system with a total of 1,500 steps of minimization, a tiny fraction of the millions of steps of MD used in the ultimate simulation of the system.

### 2.4 Solvating and Equilibrating the Membrane Model

After the complex steric clashes among the lipids have been resolved, the membrane is ready to be solvated, which presents a unique challenge to xMAS Builder. In a typical MD simulation, water simply surrounds the system of interest, and inaccuracies in the number of water molecules placed are easily accommodated by the variable simulation box size used in a constant-pressure MD simulation. In contrast, water both surrounds and is enclosed by the membranes modeled by xMAS Builder, and inaccuracies in the number of water molecules placed in the enclosed volume result in the membrane collapsing in on itself or bursting.

Because water occupies the spaces between the head groups of lipids, the number of molecules needed to solvate a membrane depends on both the composition and curvature of that membrane. The exact nature of this dependence is the subject of active investigation, but for now, we do not have a way to precisely estimate the proper number of water molecules needed to solvate a curved membrane. Thus, by necessity, xMAS Builder’s approach to solvating its membrane models involves a restrained equilibration MD simulation in which inaccuracies in the initial water molecule count reveal themselves, are quantified, and are corrected.

Running an MD simulation of the full-size model just to correct inaccuracies in the initial water molecule count would be computationally very costly, but it is fortunately possible to also equilibrate the membrane while running this simulation. Simulating the membrane while its overall shape is restrained allows the membrane’s lipids to adopt a natural packing both against one another and within the target shape of the model, and it is essential to allow this process to happen to ensure a stable final model that produces meaningful results. To set the length of the restrained equilibration simulation, we estimate that the amount of time needed to equilibrate the lipids within their local environment is approximately the same as the amount of time needed to equilibrate a membrane patch with the same target lipid composition.

#### 2.4.1 Approach

To solvate a system, xMAS Builder starts by designating a simulation box size that encloses the entire membrane, and it measures the volume enclosed by the inner surface of the membrane as well as the volume between the edges of the box and the outer surface of the membrane. Using the known density of water, xMAS Builder places the theoretically appropriate number of water molecules in both volumes, and there is then an option to add ions. Independently for the internal and external water volumes, the user specifies which ions are desired along with their concentrations, and the user specifies whether ions should be used to neutralize the membrane. xMAS Builder then calculates how many ions of each species need to be placed in each volume, and a corresponding selection of water molecules are replaced with the appropriate ions.

With the water and ions in place, xMAS Builder next sets up a restrained equilibration simulation, which gives the lipids and water a chance to adjust to the desired shape of the membrane. Using the Grid Forces feature of NAMD (introduced previously), the overall shape of the membrane is maintained, and water is prevented from passing across the membrane. Near the boundaries of the membrane, the functional form of the potential is a truncated half Gaussian, identical to what was used in the simulation to optimize the initial placement of the lipids. However, between the boundaries of the membrane, the potential is flat, meaning no forces are exerted on the core of the membrane (Fig. 5A). The form of the restraint placed on the water is similar, but the potential is flat in the bulk of the solvent, and a truncated half Gaussian acts on the solvent within the head group region of the membrane.

**Figure 5:**
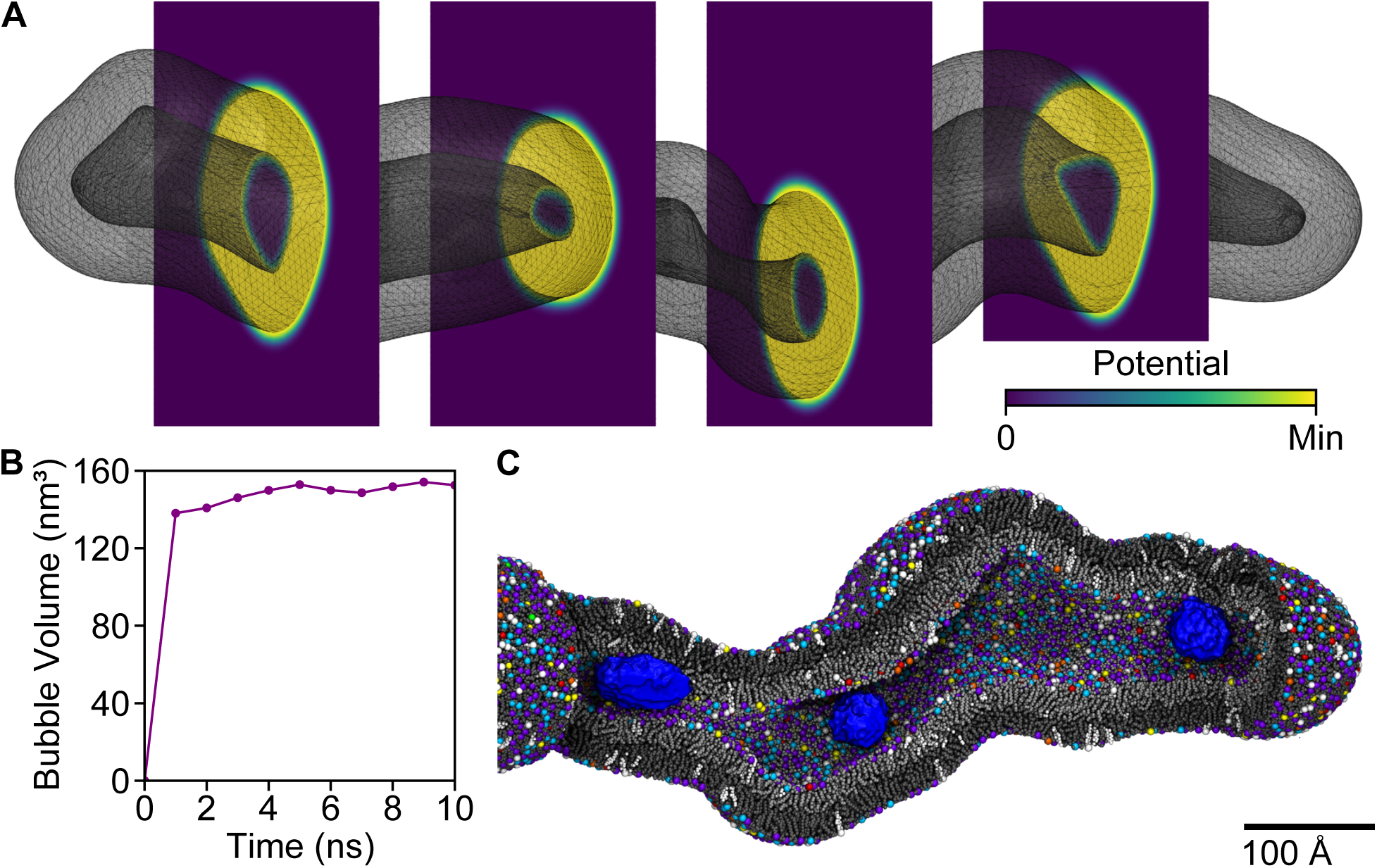
Solvating the membrane model. **(A)** Visualization of the potential used by the Grid Forces feature of NAMD to restrain the shape of the membrane during equilibration. Just outside the membrane, the functional form of the potential is a truncated half Gaussian centered on the meshes representing the membrane surfaces. Within the membrane, the potential has a uniform value. The force applied by Grid Forces is proportional to the negative gradient of the potential, so the membrane experiences no restraining force except when it pushes against the boundaries defining its target shape. **(B)** Plot of the volume of the vacuum bubbles that form in the water as a function of time. As water packs around lipid head groups in the membrane, not enough water is left to fill the volume enclosed by the membrane, and vacuum bubbles form. Once the volume of the vacuum bubbles has stabilized, xMAS Builder fills them with additional water and ions. **(C)** Cutaway molecular rendering of the vacuum bubbles (blue) that form inside the membrane model after 10 ns of equilibration. For the membrane, the same colors and molecular representations established in Fig. 1 are also used here.

As the membrane equilibrates, water molecules fill in the spaces between the head groups of the lipids, making xMAS Builders’ initial estimates of the number of molecules needed for solvation consistently low. Restrained equilibration of the system thus consistently results in vacuum bubbles forming in the water, and the volume of these bubbles is monitored as the simulation progresses. After their volume has stabilized, the bubbles are filled with water and ions, and the appropriate number of water molecules needed for this purpose can now be accurately estimated since the membrane has been fully solvated.

#### 2.4.2 Application

Using xMAS Builder, we solvated the membrane of the test system, we added 0.15 M NaCl to the water, and we performed 10 ns of restrained equilibration. The length of the simulation was determined by the ∼10 ns of simulation needed to equilibrate the planar membrane patch (as indicated by the stabilization of its thickness and side length, Fig. 2). During this time, three vacuum bubbles formed in the interior of the system, and their volume stabilized after ∼5 ns (Fig. 5). We used xMAS Builder to fill these vacuum bubbles with additional water and ions, bringing the final system size to ∼11.6 million atoms, and we then allowed the solvent to equilibrate during an additional 1 ns of restrained equilibration.

### 2.5 Simulating the Membrane Model

Following the restrained equilibration simulation, we released all restraints on the membrane and allowed it to equilibrate under unbiased conditions for 386 ns (Fig. 6). Over the course of the unrestrained simulation, the overall shape of the membrane relaxed, with areas of high local curvature becoming smoother over time. This result was expected as there was no asymmetric membrane composition or proteins present to maintain the curvature of the system. By the end of the simulation, the membrane had adopted mostly flat or semi-circular cross-sections, which makes sense as these shapes alleviate the tension caused by the unequal number of lipids in the two leaflets. Because the membrane’s global shape changed significantly over time, the simulation also tested the strength and resilience of the membrane’s construction, acting as a stress-test for xMAS Builder’s modeling protocols. While its shape relaxed, the membrane’s integrity was never compromised by any pores or fissures, indicating that the lipids were evenly distributed within the membrane and that the proper amount of water filled the model’s interior.

**Figure 6:**
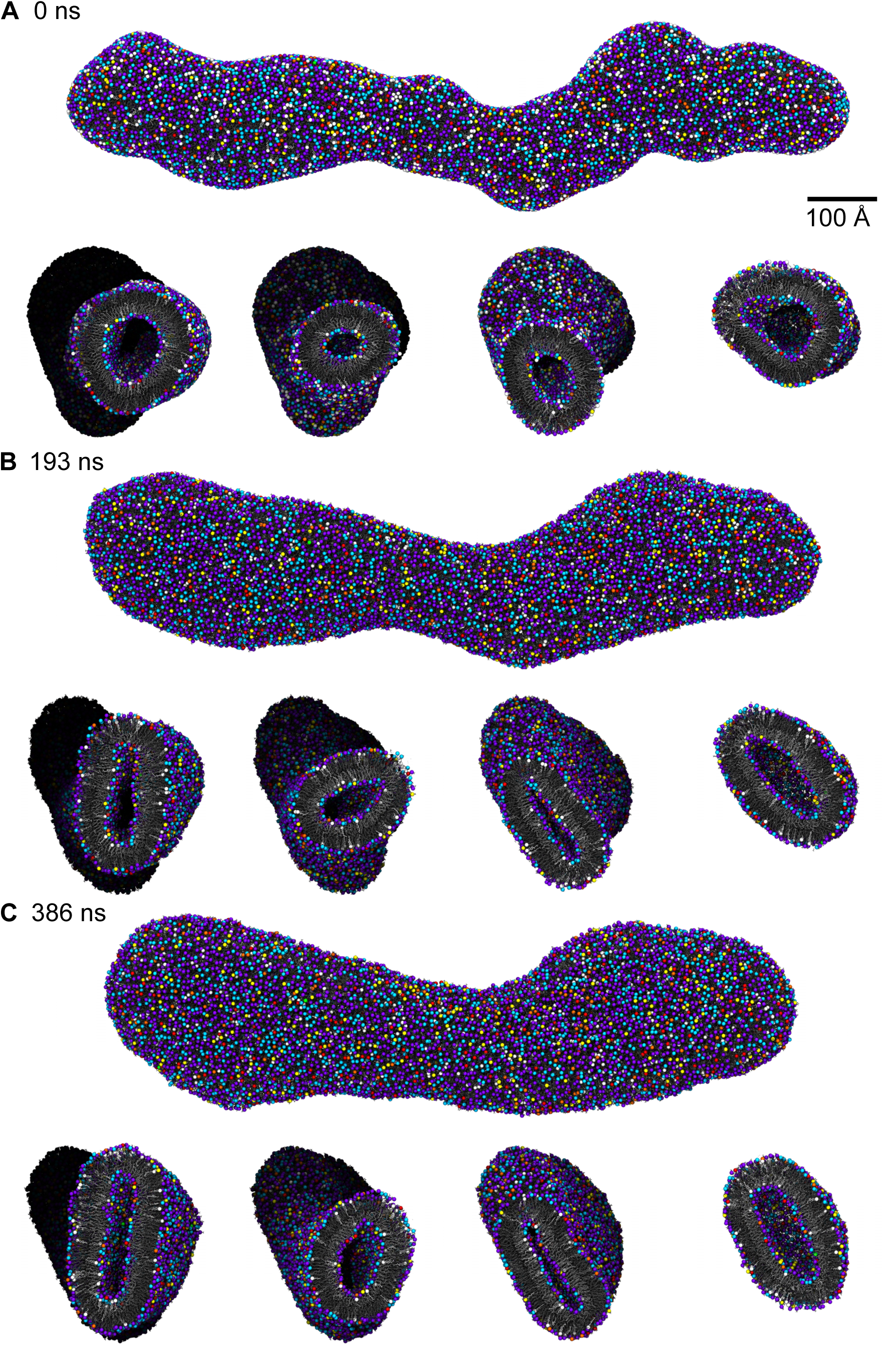
Simulating the membrane model. **(A)** Molecular renderings of the membrane after restrained equilibration. The top panel shows the whole model, and the bottom panels show cutaway renderings of the model at regular intervals along its long axis. The same colors and molecular representations established in Fig. 1 are also used here. **(B)** Molecular renderings of the membrane halfway through the simulation (after 193 ns). With no proteins or asymmetric lipid composition to enforce a specific curvature, the shape of the membrane relaxed. The fine curvature features of the initial model eroded, and the membrane flattened as much as it could. **(C)** Molecular renderings of the membrane at the end of the simulation (after 386 ns). As its shape continued to relax, the membrane lost even more of its initial complex curvature. Importantly, throughout the entire simulation, the integrity of the membrane was never compromised by pores or fissures.

### 2.6 Building and Simulating an Earlier Version of the Membrane Model

Prior to the version of the membrane model that has been discussed up to now, we used xMAS Builder to produce an earlier version of the model. The earlier model used the same shape and lipid composition and was built using the same modeling steps, but xMAS Builder used calculations based on surface area rather than volume to determine the lipid packing density (Fig. S1). This approach resulted in a model that contained 1,797 more lipids in the outer leaflet and 1,143 fewer lipids in the inner leaflet compared to the final volume-based model.

When simulated under unbiased conditions for 250 ns, the membrane’s integrity was never compromised by the formation of any pores or fissures, again illustrating the robustness of xMAS Builder’s modeling algorithms. The shape of the membrane also relaxed much like the volume-based model, with the membrane’s initial fine curvature features dissipating over time (Fig. 7). Tension caused by the unequal number of lipids in the inner and outer leaflets was eased by the model adopting mostly circular cross-sections by the end of the simulation. However, this tension was also relieved by the formation of three bilayer protrusions from the outer leaflet, which grew in size as the simulation progressed.

**Figure 7:**
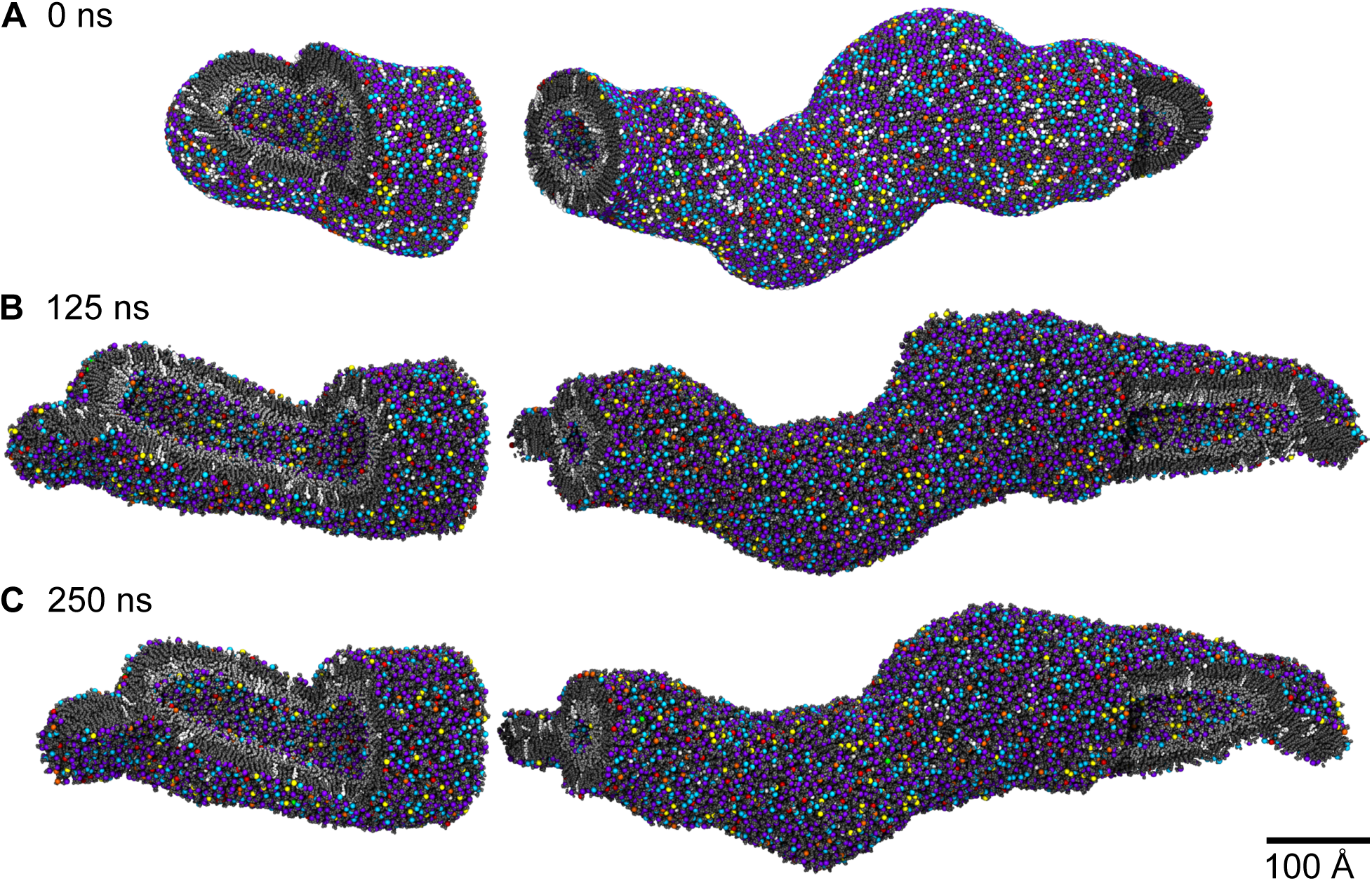
Simulating a previous version of the membrane model with a lipid packing density based on surface area. **(A)** Cutaway molecular rendering of the membrane after restrained equilibration. This version of the membrane model was built to have exactly the same shape and size as the final model, but it had 1,797 more lipids in the outer leaflet and 1,143 fewer lipids in the inner leaflet because its target packing density was determined by surface area rather than volume. The same colors and molecular representations established in Fig. 1 are also used here. **(B)** Cutaway molecular rendering of the membrane halfway through the simulation (after 125 ns). As with the volume-based model, the shape of the membrane relaxed over the course of the simulation, eroding the fine curvature features of the initial shape. Additionally, in response to overcrowding, three bilayer protrusions from the outer leaflet formed (highlighted by the thee cutaways in the rendering). **(C)** Cutaway molecular rendering of the membrane at the end of the simulation (after 250 ns). The shape of the membrane continued to relax, losing even more detail from its original shape, and the protrusions from the outer leaflet also continued to grow.

The formation of these protrusions strongly suggested to us that the outer leaflet of this model was significantly overfilled relative to the inner leaflet, and it prompted us to reevaluate our strategy to determine lipid packing density. The issue with basing packing density on surface area is that it only optimizes the packing density for the lipid head groups. In a curved membrane, this strategy leaves the lipid tails significantly over-packed or under-packed depending on the direction of the curvature, which explains why the outer leaflet was so overfilled relative to the inner leaflet in the largely tubular shape of the test system.

To address this issue, we adopted a new strategy for calculating lipid packing density based on volume, and we built a new model of the test system using it. Our intention was that this strategy would take into account the three dimensional nature of lipids and membranes, balancing slight over-packing at one end of a lipid with slight under-packing at the other. After extended equilibration of the volume-based model, there were no bilayer protrusions or any other indications of overfilled or underfilled leaflets, suggesting that this strategy accurately estimates the number of lipids needed to fill a curved membrane.

## 3 Discussion

In this work, we have developed xMAS Builder, a tool designed to build realistic atomisitic models of cellular membranes suitable for MD simulations. xMAS Builder has been designed specifically to handle arbitrarily complex lipid compositions and geometries derived from experimental lipidomics and structural (e.g., EM or ET) data. The new modeling algorithms have also been designed to scale to cellular sizes, often harnessing the existing highly developed parallel computing power in NAMD to accomplish this goal. At the same time, these new modeling algorithms (e.g., the algorithm to fix ring piercings) are also useful for traditionally sized systems.

To develop and validate the modeling algorithms of xMAS Builder, we have applied it to a test system with a representative complex geometry and lipid composition. We built two models of the test system, one containing ∼12.0 M atoms with a lipid packing density determined by surface area and one containing ∼11.6 M atoms with a lipid packing density determined by volume, and we equilibrated them with 250 ns and 386 ns, respectively, of unbiased MD. The integrity of both membranes was maintained over time while their overall shapes relaxed, but three protrusions developed on the outer leaflet of the area-based model, indicating it was overfilled relative to the inner leaflet and suggesting that volume rather than surface area should be used to determine lipid packing density. These results demonstrate that the modeling techniques developed for xMAS Builder produce high quality models, and they illustrate that these models may be used to probe the fundamental biophysics governing the structure and function of cell-scale membrane systems.

Besides xMAS Builder, TS2CG^31^ and LipidWrapper^30^ are two other tools that have also been designed to build MD-ready models of membranes with arbitrarily complex shapes. While xMAS Builder has been designed to generate membrane models for atomistic MD, TS2CG’s models are built for coarse-grained MD. The trade-offs between atomistic and coarse-grained MD simulations are well established (see,^42^ for example), and we view xMAS Builder and TS2CG as complementary tools depending on what level of resolution is desired for a particular simulation. Of course, methods are also available to change the resolution of a model from atomistic to coarse-grained and vice versa (e.g., martinize.py^43^ and backward.py^44^ from the developers of the Martini force field^45^), but we expect the current implementations of these methods will need to be adapted before they can handle cell-scale models.

Like xMAS Builder, LipidWrapper has been designed to generate models of membranes with arbitrarily complex shapes for atomistic MD, but the modeling approaches of the two tools are entirely different. In LipidWrapper’s approach, equilibrated triangular membrane patches are used to fill the input mesh’s triangles, and overlapping lipids at the triangle boundaries are simply removed from the model. This approach makes it difficult to control the lipid composition and packing density of the model, and extensive equilibration is required to heal the boundaries between the triangles and to allow the flat triangular patches to curve.^7^ LipidWrapper also has no modeling protocols to handle the unique challenge of solvating closed membrane systems. Our results indicate that proper lipid packing density and solvation have a pronounced impact on the integrity and quality of membrane models with complex closed shapes, and we suggest that xMAS Builder has an advantage over LipidWrapper in these areas.

The present implementations of the modeling algorithms used by xMAS Builder are open source and will be released upon publication of this manuscript. In future work, we plan to improve the user friendliness of these implementations with the ultimate goal of producing an easy-to-use, self-contained software package that encapsulates all of the capabilities of xMAS Builder. We also plan to continue to develop these implementations to improve their efficiency and reduce bottlenecks in xMAS Builder’s modeling protocols. Beyond improvements to the implementations of the existing modeling algorithms of xMAS Builder, we also plan to incorporate new modeling algorithms, already under development, which will allow xMAS Builder to include membrane proteins and soluble proteins in the models it builds. This additional capability will allow xMAS Builder to generate even more realistic and complex membrane models, enabling detailed characterization of the structural dynamics of these systems with MD simulations.

## 4 Methods

Unless otherwise specified, all MD simulations were performed with NAMD^34^ using the CHARMM36m force field^36–39^ at a constant temperature of 310 K and a constant pressure of 1 atm. Constant temperature was maintained using Langevin dynamics for all non-hydrogen atoms with a damping coefficient of 1 ps^−1^. Constant pressure was maintained using a Nosé-Hoover Langevin piston^46, 47^ with a period of 100 fs and a damping time scale of 50 fs. Simulations were performed with an integration time step of 2 fs. The cut-off distance for both electrostatic and van der Waals interactions was set to 12 Å, and a switching function was applied at 10 Å. Periodic boundary conditions were applied in all simulations, and long-range electrostatic interactions were calculated using the particle mesh Ewald method^48, 49^ with a grid point density of 1 Å^−1^.

## 5 Data Availability

All data generated or analyzed in this study are included in this manuscript and its supplementary information files.

## 6 Code Availability

The source code for xMAS Builder will be released upon publication of this manuscript.

## 7 Acknowledgments

Research reported in this publication was supported by grants from the National Institutes of Health under award numbers P41-GM104601 and R24-GM145965. NT acknowledges support from the National Science Foundation Graduate Research Fellowship Program under Grant No. 1746047. Any opinions, findings, and conclusions or recommendations expressed in this work are those of the authors and do not necessarily reflect the views of the National Institutes of Health or the National Science Foundation. This work used resources of the Extreme Science and Engineering Discovery Environment (XSEDE), which was supported by National Science Foundation grant number ACI-1548562, through allocations TG-MCA06N06 and TG-MCA93S028. This work also used resources of the Oak Ridge Leadership Computing Facility, which is a Department of Energy Office of Science User Facility supported under Contract DE-AC05-00OR22725, through Innovative and Novel Computational Impact on Theory and Experiment (INCITE) allocations BIP117 and BIP115 and Summit Early Science Program (ESP) allocation BIP177. This work finally used resources from Blue Waters, which was supported by the National Science Foundation (awards OCI-0725070 and ACI-1238993) and the State of Illinois, through Petascale Computing Resource Allocations (PRAC) project BANV. The authors gratefully acknowledge helpful discussions with John Stone, Jim Phillips, Dave Hardy, Chris Mayne, and Josh Vermaas.

## 8 Author Contributions

ET conceived and supervised the study, and NT designed and executed it. NT prepared the initial draft of the manuscript, and ET and NT revised it together.

