## Supplementary material for "Enabling Atomistic Modeling and Simulation of Complex Curved Cellular Membranes with xMAS Builder": Figure S1

**A**

Outer Leaflet

$$605{,}837\ \text{Å}^2 \bullet \frac{1{,}000\ \text{lipids}}{52{,}441\ \text{Å}^2} = 11{,}552\ \text{lipids}$$

Inner Leaflet

$$272{,}799\ \text{Å}^2 \bullet \frac{1{,}000\ \text{lipids}}{52{,}441\ \text{Å}^2} = 5{,}202\ \text{lipids}$$

**B**

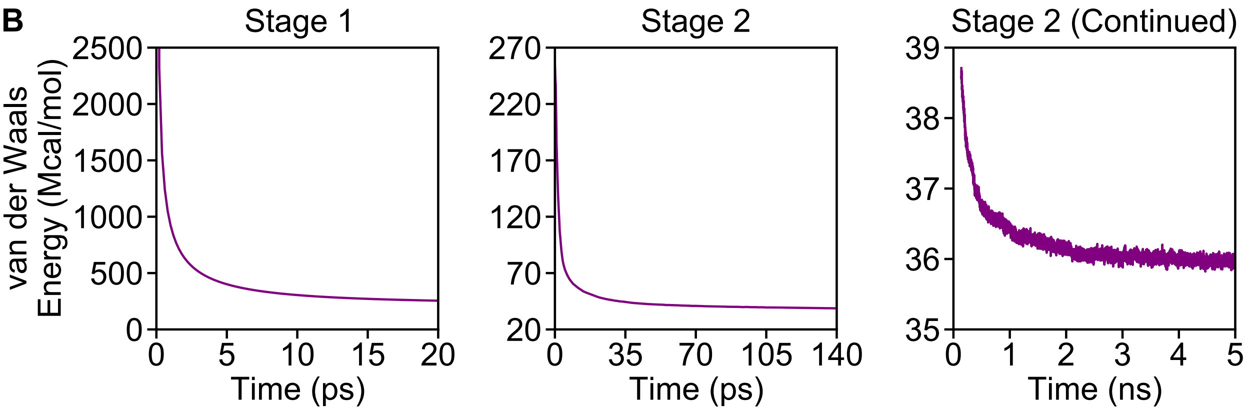


**C**

| | |
|---|---|
| Total Lipids: | 16,754 |
| Ringed Lipids: | 3,111 |
| Ring Piercings: | 880 → 12 → 2 → 1 → 0 |
| "Frozen" Lipids: | 517 → 7 → 1 → 1 → 0 |
| Entanglements: | 6 → 0 |

**D**

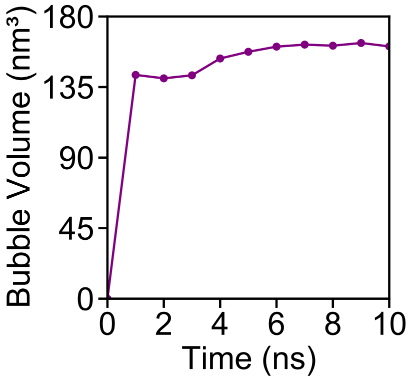


**Figure S1: Building a previous version of the membrane model using a lipid packing density based on surface area. (A)** Calculations of the number of lipids used to fill the inner and outer leaflets in the model. In these calculations, the area of the meshes representing the inner and outer surfaces of the membrane are multiplied by the lipid packing density of the equilibrated membrane patch (Fig. **??**). **(B)** Plots of the total particle interaction energy during the simulation to optimize the initial positions of the lipids. As with the later version of the model, this simulation occurred in two stages, with the first stage (20 ps) utilizing velocity quenching, and the the second stage (5 ns) proceeding without it. **(C)** Tabulation of the number of ring piercings and lipid entanglements corrected by xMAS Builder for this version of the model. **(D)** Plot of the volume of the vacuum bubbles that formed in the water during restrained equilibration. As with the later model, the volume stabilized after ∼5 ns and were subsequently filled with additional water and ions.
